# Chronic Cognitive and Cerebrovascular Function Following Mild Traumatic Brain Injury in Rats

**DOI:** 10.1101/2022.01.05.474992

**Authors:** Daniel R. Griffiths, L. Matthew Law, Conor Young, Alberto Fuentes, Seth Truran, Nina Karamanova, Laura C. Bell, Gregory Turner, Hannah Emerson, Diego Mastroeni, Rayna Gonzales, Peter D. Reaven, Chad C. Quarles, Raymond Q. Migrino, Jonathan Lifshitz

## Abstract

Severe traumatic brain injury results in cognitive dysfunction in part due to vascular perturbations. In contrast, the long-term vasculo-cognitive pathophysiology of mild TBI (mTBI) remains unknown. We evaluated mTBI effects on chronic cognitive and cerebrovascular function and assessed their interrelationships. Sprague-Dawley rats received midline fluid percussion injury (N=20) or sham (N=21). Cognitive function was assessed (3- and 6-month novel object recognition (NOR), novel object location (NOL) and temporal order object recognition (TOR)). 6-month cerebral blood flow (CBF) and blood volume (CBV) using contrast MRI and *ex vivo* pial artery endothelial and smooth muscle-dependent function were measured. mTBI rats showed impaired NOR, with similar (non-significant) trends in NOL/TOR. Regional CBF and CBV were similar in sham and mTBI. NOR correlated with CBF in lateral hippocampus, medial hippocampus and primary somatosensory barrel cortex while inversely correlating with arterial smooth muscle-dependent dilation. 6-month baseline endothelial and smooth muscle-dependent arterial function were similar among mTBI and sham, but post-angiotensin II stimulation, mTBI showed no change in smooth muscle-dependent dilation from baseline response, unlike the reduction in sham. mTBI led to chronic cognitive dysfunction and altered angiotensin II-stimulated smooth muscle-dependent vasoreactivity, a paradigm that could advance understanding of the long-term sequelae of human mild TBI.

## INTRODUCTION

It is estimated that 61 million individuals worldwide experience traumatic brain injury (TBI) from all causes every year.^1, 2^ Nearly 20% of the more than 2.6 million US service members deployed to Operation Enduring Freedom and Operation Iraqi Freedom have sustained at least one TBI event.^3^ TBI may result in lifelong disability and survivors can face enduring motor, cognitive and social impairments.^4^ Secondary brain injury following TBI is caused by a combination of neuronal and vascular damage, proteolytic pathways, free radical damage, apoptosis and inflammatory processes.^5^ Cerebrovascular dysfunction plays an important role in severe, acute TBI with ischemic brain damage evident at autopsy in >90% of acute TBI mortalities.^6^ The lifetime consequences of TBI include long-term cognitive dysfunction, which may be associated with chronic traumatic encephalopathy.^7, 8^

Unlike repetitive and severe TBI, the long-term cognitive and vascular function in mild TBI (mTBI) remain poorly characterized. mTBI is the most common form of TBI among civilians and military service members, occurring in ~82% of military TBI.^9^ Epidemiologic data indicate that individuals with TBI history have a higher risk of developing dementia.^10–12^ Among World War II Navy and Marine veterans, nonpenetrating head injury in early adulthood was associated with increased risk of Alzheimer’s disease (AD) and other dementias.^13^ Of interest, even mild head injury was found to be a predisposing factor for AD or dementia.^10, 14^ A longitudinal study of human mTBI (post-concussion) show impaired cerebral blood flow (CBF) in the acute (1 day and 1 week) setting that closely correlated with neuropsychiatric symptoms, and global CBF recovered by 1 month.^15^ The relationship between vascular and cognitive function was further supported by their finding that persistent impaired CBF at 1 month in the dorsal midinsular cortex was associated with slow recovery of neuropsychiatric symptoms. In separate studies, experimental severe TBI led to cognitive dysfunction^16–19^ and impaired CBF^20^ at 1-year post-injury; similar preclinical studies on chronic mTBI are not available. Thus, vascular dysfunction is likely related to long-term cognitive deficits, and therefore co-exist in chronic mTBI, but empiric evidence of this relationship is still lacking.

Experimental models of mTBI provide a unique opportunity to investigate chronic effects as well as relationships between cognitive and cerebrovascular function. Midline fluid percussion injury (FPI) is a well-validated model of mTBI.^21^ Mechanical forces of the fluid pulse reflect off the temporal ridge of the skull to primarily affect the hippocampal area CA3, primary somatosensory barrel cortex (S1BF) and ventral posterior nuclei of the thalamus. The model relates to non-catastrophic TBI with acute physiologic disruption that recovers within a few days with absence of gross histopathologic damage and lack of cavitation months post-injury.^22, 23^

The aims of this study are to determine the chronic effects of mTBI on cognitive and cerebrovascular function and to evaluate the relationship between the two.

## METHODS

### Animals

The study was approved and supervised by the University of Arizona Institutional Animal Care and Use Committee. Male Sprague Dawley rats (~9-10 weeks old, ~300g-325g, Charles River Labs) were given one week to acclimate in their home cages. Rats were given standard chow and water *ad libitum* and were housed in a reverse light cycle room. Experiments were conducted in accordance with University of Arizona, Department of Defense Instruction 3216.01, RIGOR and Animal Research: Reporting In Vivo Experiments (ARRIVE) guidelines concerning the care and use of laboratory animals. Adequate measures were taken to minimize pain or discomfort.

### Midline Fluid Percussion Injury

All rodent experiments were conducted in cohorts of uninjured (sham) and mTBI animals, randomly assigned to groups at the time of brain injury. The individual inducing the brain injury was different from those assessing cognitive function, *in vivo* imaging data, or *ex vivo* vascular function. After data collection, the group assignment for each animal was decoded. Rats were subjected to midline FPI similar to methods described previously.^21, 22, 24^ The midline variation of FPI represents mild clinical brain injury because of the acute transient behavioral deficits, the late onset of behavioral morbidities, and the absence of gross histopathology.^22^ Briefly, rats were anesthetized with isoflurane for surgery, but did not receive systemic analgesia before injury. Body temperature was maintained with a Deltaphase^®^ isothermal heating pad (Braintree Scientific Inc., Braintree, MA). In a head holder assembly (Kopf Instrument, Tujunga, CA), a midline scalp incision exposed the skull. A 4.8-mm circular craniotomy was performed (centered on the sagittal suture midway between bregma and lambda) without disrupting the underlying tissue. An injury cap was fabricated from a Luer-Lock needle hub, which was cut, beveled, and scored to fit within the craniotomy. A skull screw was secured in a 1-mm hand-drilled hole into the right frontal bone. The injury hub was affixed over the craniotomy using cyanoacrylate gel and methyl-methacrylate (Hygenic Corp., Akron, OH) was applied around the injury hub and screw. The incision was sutured at the anterior and posterior edges and topical bacitracin and lidocaine ointment were applied. Animals were returned to a warmed holding cage and monitored until ambulatory (approximately 60-90 min).

For injury induction, animals were re-anesthetized with isoflurane. The dura was inspected through the injury-hub assembly, which was then filled with normal saline and attached to the fluid percussion device (Custom Design and Fabrication, Virginia Commonwealth University, Richmond, VA). Animals were randomly assigned to receive brain injury (N=20; 2.29 ±0.02 atm, 2.20-2.44 atm) or sham injury (N=21) by releasing (or not releasing) the pendulum onto the fluid-filled cylinder. Animals were monitored for the presence of a forearm fencing response and the return of the righting reflex as indicators of injury severity.^24^ After injury, the injury hub assembly was removed *en bloc*, integrity of the dura was observed, bleeding was controlled with saline and gauze, and the incision was stapled. Brain-injured animals had righting reflex recovery times of 522±34 (range 245-755) seconds and sham-injured animals recovered within 15 seconds. After recovery of the righting reflex, animals were placed in a warmed holding cage before being returned to the housing room. Each rat was evaluated for post-operative health for three days. Appropriate interventions, including but not limited to injections of subcutaneous bolus of saline, and feeding with a wet mash of food and water, were performed if clinically indicated. Rats were euthanized if their body weights fell below 10% of their own pre-operative body weight; none met these criteria.

### Cognitive Function Evaluation

The assessments were performed by investigators blinded to treatment allocation at 3 and 6 months post-injury. Object recognition tasks take place in a square arena (68.58 cm x 68.58 cm) (Figure 1A-B). White noise was used to mask environmental noises. The novel object recognition task (NOR) tested short-term recognition memory.^25^ Rats acclimate to the arena for 3 minutes. Rats are then presented with two objects (O1, O2) in opposite corners of the arena (5 minutes) in the sample trial. After a 4-hour delay, rats are returned to the arena with one object (O2) being replaced by a novel object (O3). Normal rats explore the novel object (O3) more than the familiar object.^25^ The novel object location task (NOL) tested long-term spatial memory.^26^ The test trial of the NOR serves as the sample trial. After 24 hours, O3 was placed in the same place as the previous day and O1 was moved to an adjacent corner in the arena. Normal rats explore the item in the novel location (O1) more than the unmoved object. The temporal order object recognition task (TOR) tested temporal working memory by the ability to recognize the order of objects presented over time.^26^ Two sample trials established the cognitive framework, followed by a test trial. In sample trials, rats explored 2 copies of an object for 5 min, followed by exploration of a separate pair of identical objects followed by the test trial where one of each item was present. Breaks between sample phases were 3 minutes before the two sample phases and 5 minutes before the test trial (5 min). Normal rats explore the initial object, rather than the more recent object.

**Figure 1.**
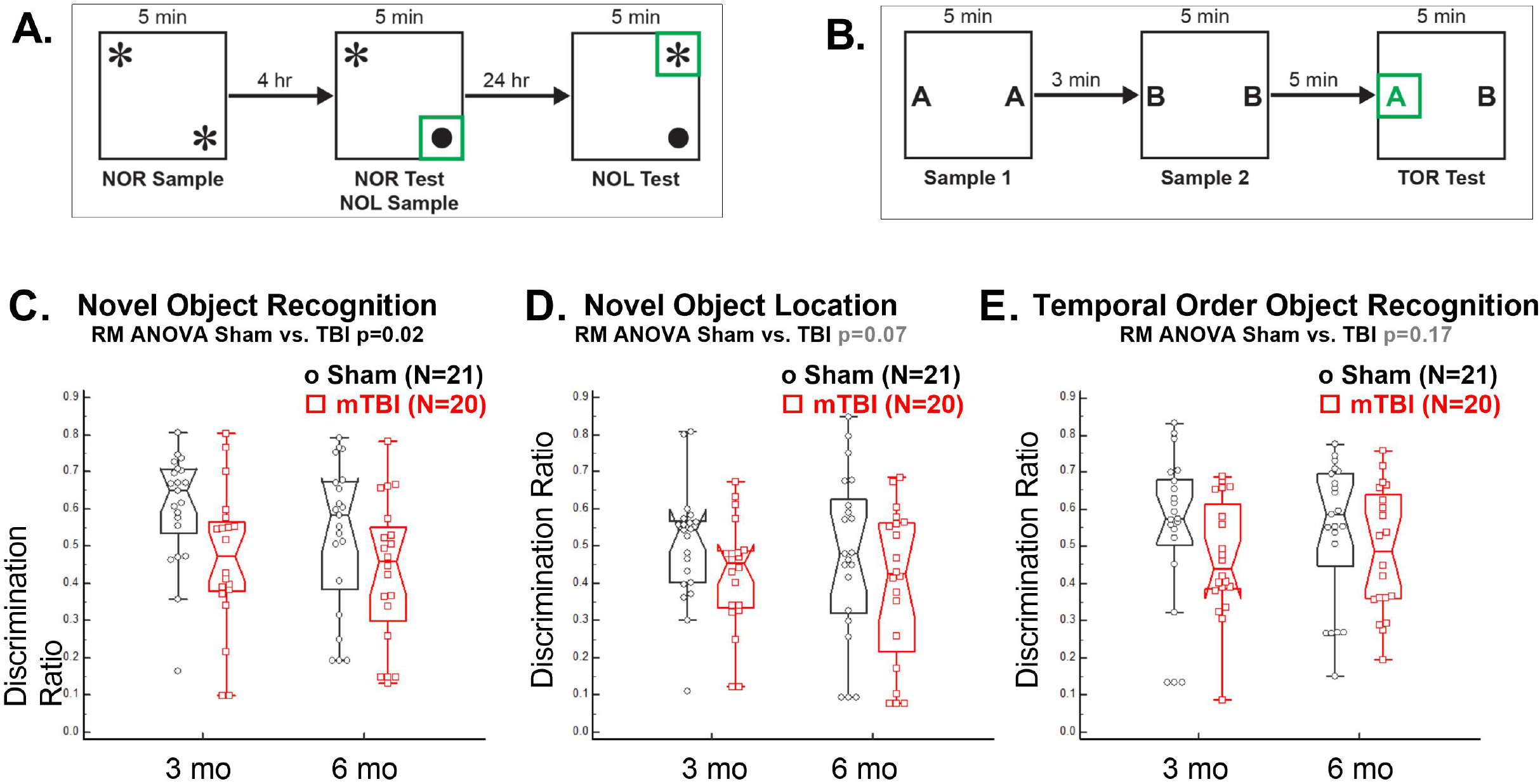
Chronic cognitive impairment following mTBI. A. Schematic of object recognition tasks. Novel object recognition (NOR) tests short term memory by replacing an object (*) with (•) after a 4-hour delay. Novel object location (NOL) tests long term memory by shifting the position of the familiar object (*) after a 24-hour delay. B. Temporal order object recognition (TOR) tests working memory by presenting pairs of objects. C. There was impaired novel object recognition at 3 and 6 months in rats subjected to mTBI versus sham controls. D-E. There were similar trends but not statistically significant differences in novel object location and temporal order object recognition between mTBI and sham rats.

Exploration of an object was defined as the nose being within ~2 cm of the object. Differences in time spent exploring each object were recorded for all tasks and used to create a discrimination ratio (exploration time of the target object/exploration time of both objects). Normal rats explore the target object more than the original object, in this case resulting in discrimination ratio above 0.5.^26^ A discrimination ratio of 0.5 indicates equal exploration of object, and equivalent to chance performance. Every trial was tracked and recorded using Ethovision software (Noldus, Leesburg, VA).

Rats were required to explore each item more than 5 seconds during the sample trial(s). If a rat did not meet exploration criteria, a discrimination ratio was imputed (Supplement Table 1). The imputed minimal discrimination ratio was calculated using the average lowest exploration time for the target object and the highest average exploration time for the specific rat’s experimental condition.

### Magnetic Resonance Imaging

Rats underwent brain magnetic resonance imaging (MRI) 6 months post-injury. Scanning was performed using a 7T small animal, 30-cm horizontal-bore magnet and BioSpec Avance III spectrometer (Bruker, Billerica, MA) with a 116 mm high power gradient set (600 mT/m) and either a 70mm rat volume quadrature coil or 40mm rat head volume quadrature coil depending size and weight of the animal. Each animal was anesthetized and maintained under isoflurane anesthesia (1-2%) in medical air. A tail vein catheter was place for injection of ferumoxytol (FERAHEME) at a dosing scheme of 0.3 mg/kg for first pass injection at 2.2mL/min and 0.7mg/kg for slow infusion at 1.1 mL/min using a Genie Touch (Kent Scientific: Torrington, CT) power injector. Respiration was continually monitored via a pillow sensor positioned under the abdomen (SA Instruments, Stoney Brook, NY). Normal body temperature (36-37°C) was maintained with a circulating warm water blanket (Thermo Scientific, Rockford, IL).

Anatomic T2-weighted images were acquired with RARE sequence (TR:5500 TE: 12.5 FA: 90°, Avgs: 4, FOV: 30×30mm, MTX: 150×150, Slices: 50, Slick Thickness: 0.5mm) for registration and localizing hippocampus. Relaxometry measurements were acquired with pre/post contrast T1 maps with a RAREVTR sequence (TE: 8.5, TR: [275 500 800 1100 2000 4000], FA: 90, RARE Factor: 4, Avgs: 2, FOV: 30×30mm, MTX:150×150, Slices: 6, Slice Thickness: 1mm) and T2* maps with multi-gradient-echo (MGE) sequence (TR: 1000, FA: 45, Avgs: 6, TE: [4 8 12 16 20 24], FOV: 30×30mm, MTX: 150×150, Slices: 6, Slice Thickness: 1). First Pass Imaging utilized an EPI sequence (TE: 7.5, FA: 45,TR: 1000, Avgs:1,FOV: 30×30mm, MTX:64×64, Slices:6, Slice Thickness: 1mm, Reps: 240) allowing for 1 minute of baseline image acquisition before injection and 3 minutes of post-injection image acquisition.

### Magnetic Resonance Imaging Analysis

The perfusion datasets were analyzed by investigators blind to treatment allocation using in-house code developed in MATLAB and the MATLAB Imaging Processing toolbox for registration functions (Mathworks, Natick, Massachusetts). Pre-processing steps included rigid registration and arterial input function (AIF) determination. The first time point of the DSC dataset was registered to the anatomical T2 images in order to apply the tissue ROIs. The remaining DSC time points were registered across time to account for any potential movement during the scan. After registration, ΔR2* time curves were computed using the conventional single-echo equation.^27^ The AIF was automatically determined using a previously published algorithm.^28^ Mean tissue curves were found in each of the pre-determined ROIs within the brain as mentioned above. Finally, CBF was determined as the maximum value of the residue function – the end product of the deconvolution.^29^ To compensate for any potential delay between the time curves, the AIF was discretized using a block-circulant method prior to the deconvolution and signal-to-noise (SNR) based thresholding was used truncate the inverse matrix to help regularize this discretized AIF.^30^ CBF and CBV were calculated from coronal slices. Regional measurements were made for S1BF and the medial and lateral hippocampus (bisected medial to the dentate gyrus). The length and width of the third ventricle (~3 mm posterior from bregma) were measured and compared to assess for ventriculomegaly. Evidence of gross bleeding and/or microhemorrhage was evaluated by gross visualization from T2*-weighted or susceptibility images.

### Ex-vivo cerebrovascular function assessment

Rats were euthanized at 6 months post-injury via sodium pentobarbital overdose and pial arteries (circle of Willis) were carefully dissected from the brain and placed immediately in HEPES buffer. The methods were adapted from previous work.^31, 32^ Middle or posterior cerebral arteries were isolated and cannulated, and vessel luminal diameters were measured using videomicroscopes throughout the procedure by investigators blind as to treatment allocation. The vessels were pressurized to 30 mm Hg for 30 minutes and then 60 mm Hg for 30 minutes. Myogenic tone was measured based on dilator response to intraluminal pressure. For each rat, several arterial segments (3-4) were cannulated and underwent vasoreactivity measurements at baseline and following 1-hour exposure to 1 of 3 vascular agonists. Following stabilization, each vessel was preconstricted with increasing doses of endothelin-1 (10^-9^-10^-4^M, Sigma Aldrich, St. Louis MO) until ~60% of last observed maximum diameter was achieved. Baseline dilator responses to acetylcholine (increasing doses from 10^-9^-10^-4^M) were measured to assess endothelium-dependent vasodilation, followed by 10^-4^M of nitric oxide donor diethylenetriamine NONOate (DETA NONOate, Cayman Chemicals, Ann Arbor MI) to evaluate smooth muscle-dependent dilation. To assess arterial response to vascular agonists/stressors, baseline (control) response was followed by washout, and each artery was assigned to exposure for 1 hour to one of the following treatments: angiotensin II (20 μM, Sigma Aldrich), β-amyloid 1-42 (Aβ42, 1 μM, Anaspec, Fremont CA)^31, 33^ or high glucose (33 mM)^34^ and a second measurement of dilator responses to acetylcholine and DETA NONOate was performed. For the arterial segments assigned to angiotensin II, baseline intraluminal pressure prior to vasoreactivity experiments was increased from 0, 30, 60 and then to 90 mm Hg. Post-treatment dilator responses were compared to baseline control response in both sham and mTBI rats and change in dilator responses (treatment minus baseline) was compared between sham and mTBI rats.

### Data and Statistical Analyses

Sample size considerations: Both cerebrovascular and cognitive function outcomes represent study primary outcomes. Our separate preliminary pilot study (N=3) on pial arterial function 5 weeks following surgery showed a difference in dilator response to 10^-4^ M acetylcholine between mTBI and sham of −13.8% with a combined standard deviation of 8.9%. A sample size of at least N=15 per group would allow us to show a similar difference at 6 months post-surgery but with a more conservative standard deviation of 13% (α=0.05, β=0.80).

Cognitive function measures (NOR, NOL, TOR) were analyzed using repeated measures analysis of variance (ANOVA) (two-factor study with repeated measures on one factor) with time (3- and 6-month) as within-subject factor and treatment (mTBI, sham) as grouping/between-subjects factor. mTBI and sham MRI and ex-vivo vasoreactivity data were compared using unpaired t-test for normally distributed data or Mann-Whitney test for data that were not normally distributed. Shapiro-Wilk test was used to determine normality of distribution. Comparison of dilator responses before and following exposure to vascular agonists was done using paired t-test. Correlation analyses were performed using Pearson’s method for normally distributed data or Spearman’s method for non-normally distributed data. Significant p value was set at p<0.05 (2-sided). Analyses were performed using MedCalc version 19.8 (MedCalc Software, Ostend Belgium). Data in Figures 1 and 2 are represented as individual datapoints and as notched boxplot format (showing median, interquartile range, notch representing 95% confidence interval of the median, upper whisker as lesser of 75th percentile or maximum value and lower whisker as greater of 25^th^ percentile or minimum value). Preclinical data can be made available after official request to the corresponding author.

**Figure 2.**
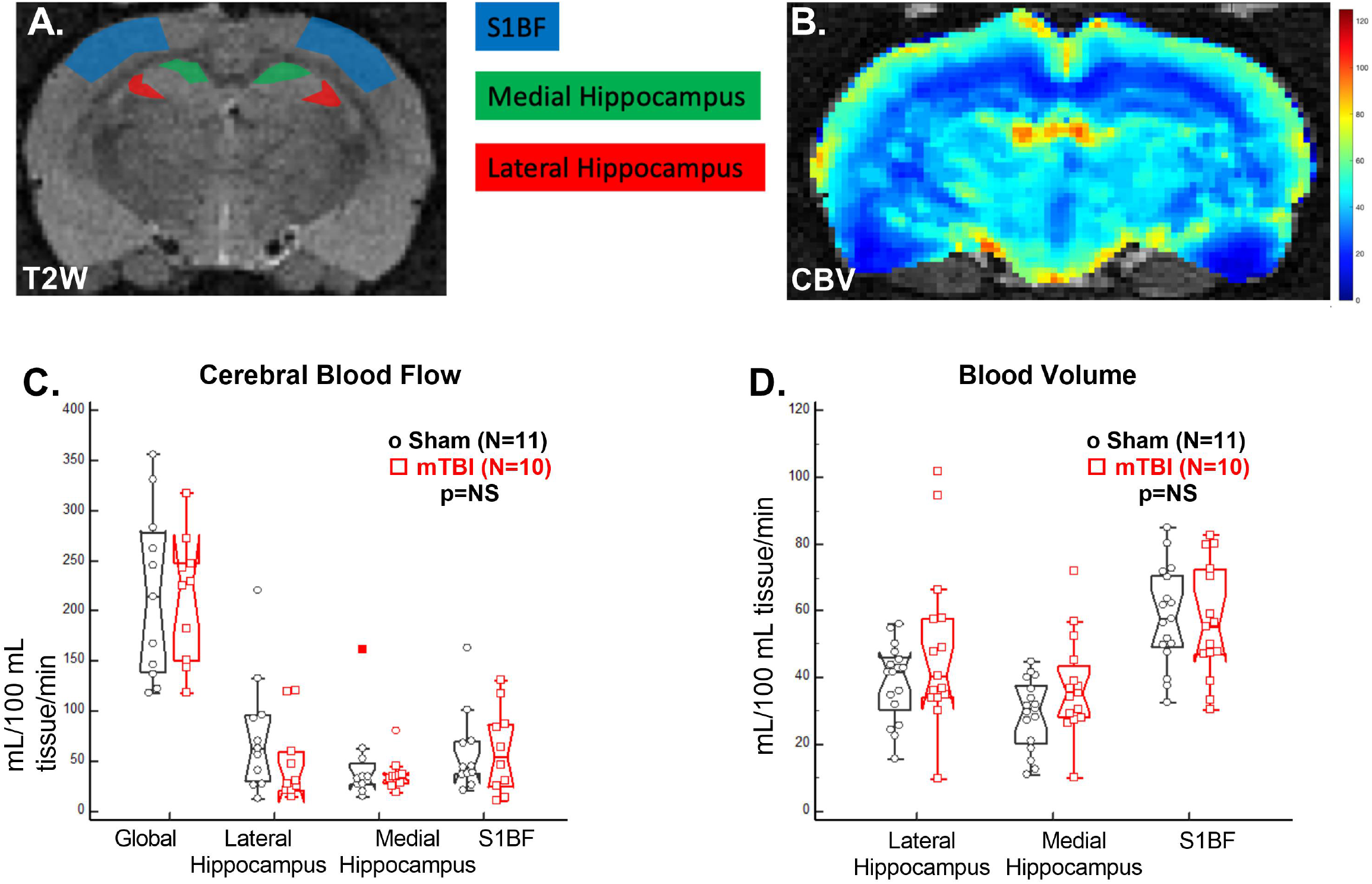
In vivo resting cerebral perfusion. A. Representative T2-weighted anatomic image. The colored areas represent regions of measurement. B. Parametric regional cerebral blood volume map. C. Global and regional resting cerebral blood flow did not differ between mTBI and sham rats at 6 months. D. Regional resting blood volume also did not differ between mTBI and sham rats.

## RESULTS

### mTBI Leads to Persistent Cognitive Impairment

Repeated measures ANOVA showed significant difference by treatment group factor (mTBI versus sham) in NOR discrimination ratio (p=0.02), but not by time factor (3-versus 6-months) (p=0.09) (Figure 1C). The NOR results indicate persistent impairment in cognitive function relating to short-term recognition memory^25^ following mTBI. A similar trend by treatment group factor was seen in NOL and TOR, although the differences did not reach statistical significance (Figure 1D-E).

### Effects of mTBI on Chronic Resting CBF and CBV

Representative T2-weighted MRI brain structural image (Figure 2A) and parametric mapping of CBV (Figure 2B) are shown, including delineation of brain regions analyzed. There was no significant difference in resting global or regional CBF between mTBI and sham rats at 6 months (Figure 2C). There was also no difference in regional CBV between mTBI and sham (Figure 2D). There was no gross bleeding and/or micro hemorrhage noted among sham and mTBI rats and there were no significant differences in length (sham: 0.28±0.004; mTBI: 0.27±0.01 mm, p=NS) or width (sham: 0.03±0.0.002; mTBI: 0.03±0.004 mm, p=NS) of the third ventricle to suggest ventriculomegaly. Post-mortem histopathology confirmed no evidence of blood extravasation in subcortical white matter (data not shown).

### Association Between CBF and Cognitive Function

There were significant correlations between 6-month NOR and the following vascular parameters: lateral hippocampus CBF, medial hippocampus CBF and S1BF CBF (Figure 3A-C and Supplement Table 2). There was a significant correlation between 6-month NOL and medial hippocampus CBF (Figure 3D and Supplement Table 2). The results demonstrate the association between cognitive function and resting regional cerebrovascular perfusion 6 months following mTBI or sham procedure.

**Figure 3.**
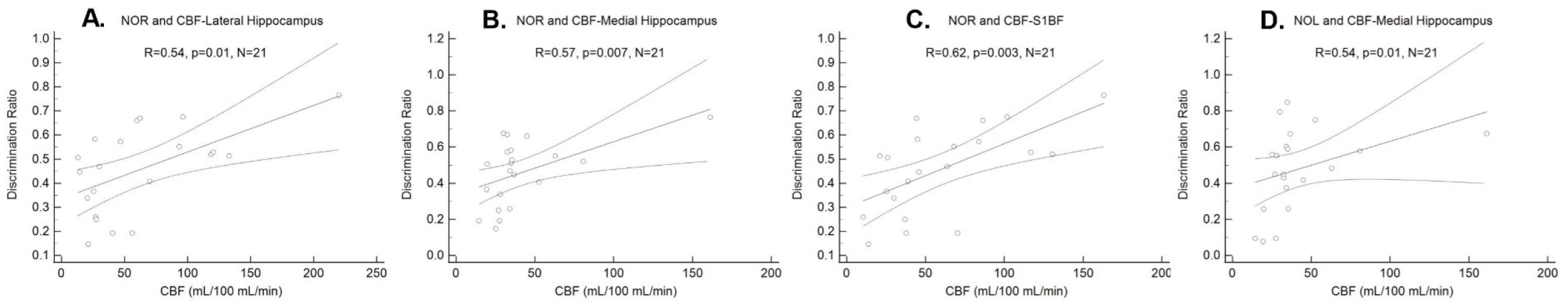
Relationship between cognitive and vascular function.6-month NOR was directly associated with CBF in the lateral hippocampus (A), medial hippocampus (B) and S1BF (C). 6-month NOL was directly associated with CBF in medial hippocampus. Data are from sham and mTBI rats.

### Effects of mTBI on Pial Cerebral Arterial Function

Constriction responses to increasing intraluminal pressure in ex-vivo pial cerebral arteries did not differ between mTBI and sham rats (Figure 4A-B), signifying lack of difference in myogenic tone. There were also no differences in baseline (unstimulated) dilator response between mTBI and sham rats when arteries were exposed to increasing doses of acetylcholine, signifying lack of difference in baseline endothelium-dependent function, or when arteries were exposed to DETA NONOate, signifying lack of difference in baseline smooth muscle dependent function (Figure 4C).

**Figure 4.**
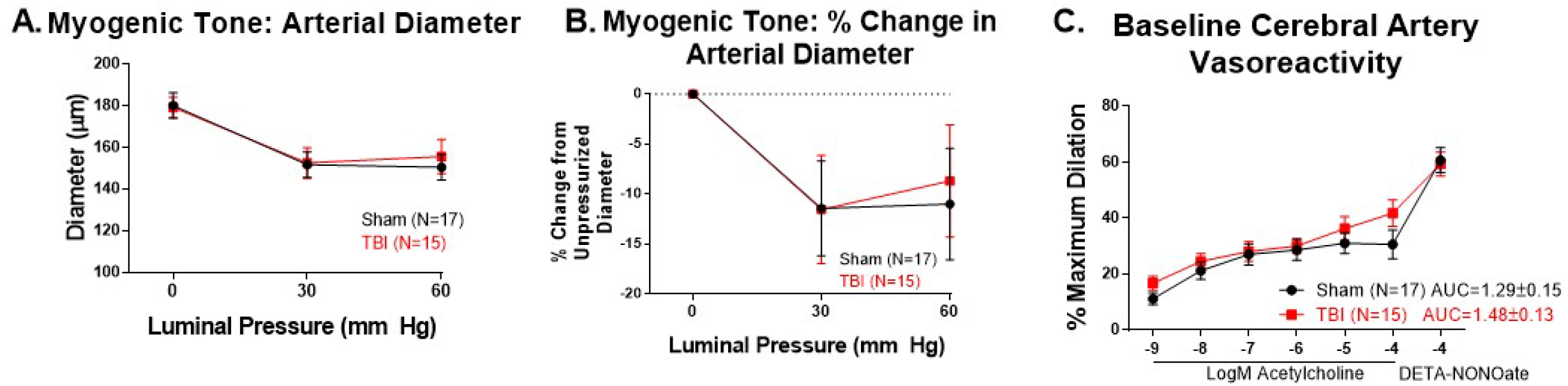
Myogenic tone and baseline vasoreactivity. A-B. There was no significant difference in response to increasing intraluminal pressure between TBI and sham rat cerebral arteries. % change in arterial diameter was calculated as (Diameter30 or 60 - Diameter0)/Diameter0 x 100%) C. Dilator responses to increasing doses of acetylcholine and DETA-NONOate were also not significantly different between TBI and sham rats.

Dilator responses to acetylcholine and DETA-NONOate were compared at baseline (control) and following exposure to vascular agonists in sham and mTBI cerebral arteries. There was significant reduction in dilator response to acetylcholine and DETA-NONOate following exposure to angiotensin II in sham rats (Fig. 5A1) but not in mTBI rats (Fig. 5A2). The change in dilator response to DETA-NONOate following angiotensin II exposure was significantly different between sham and mTBI with reduction in sham but not with mTBI, suggesting absence of smooth muscle constriction response following angiotensin II exposure in mTBI versus sham. Based on area under the curve (representing combined response to increasing acetylcholine doses), exposure to high glucose showed no difference with baseline response in both sham and mTBI (Figure 5B1-2), but HG resulted in significant decrease in dilator response to DETA-NONOate in both sham and mTBI, with no significant difference in response between sham and mTBI (Figure 5B3). Dilator response to acetylcholine was marginally reduced in sham following Aβ42 but not in mTBI (Figure 5C1-2) but dilation to DETA-NONOate was reduced in both sham and mTBI. The Aβ42-induced change in dilator response was not different between sham and mTBI (Figure 5C3). Overall, these results suggest reduced vasoconstrictor response post-angiotensin II exposure in mTBI versus sham cerebral arteries, but no difference between sham and mTBI with high glucose or Aβ42.

**Figure 5.**
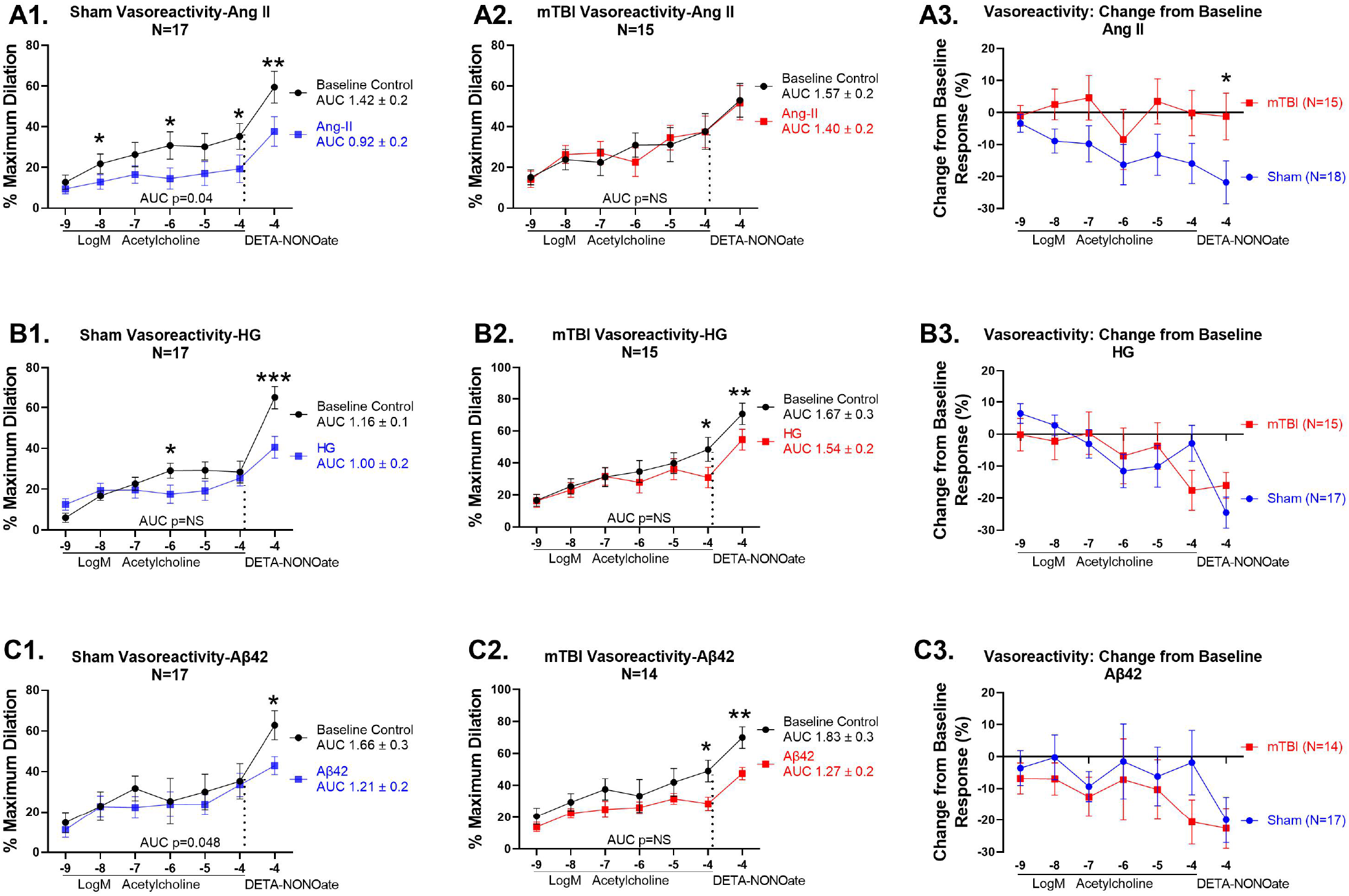
Pial artery dilator responses at 6 months following exposure to vascular agonists. A. Sham rats showed reduced dilator response to acetylcholine and DETA-NONOate following exposure to angiotensin II (A1), which was not seen in mTBI rats (A2). There was significant reduction in dilation to DETA-NONOate in sham compared to mTBI (A3). B. Overall dilator response to acetylcholine (AUC) was not different following high glucose exposure in both sham and mTBI, while dilation to DETA-NONOate was reduced in both sham and mTBI (B1-B2). There was no difference in change in dilator response to high glucose between sham and mTBI (B3). C. There was marginal reduction in overall dilator response to acetylcholine (AUC) following Aβ42 in sham but not in mTBI, but dilation to DETA-NONOate was significantly reduced in both sham and mTBI (C1-2). There was no difference in change in dilator response to Aβ42 between sham and mTBI (C3). *p<0.05, **p<0.01, Ang II-angiotensin II, HG-high glucose

### Association of ex vivo pial arterial vasoreactivity with cognitive function and in vivo regional blood volume

Pial arterial dilator response to DETA NONOate was inversely related to 6-month NOR (Figure 6A, Supplement Table 3). These data show the association between cerebral arterial smooth muscle function and cognitive function. There was no significant correlation between change in post- and pre-angiotensin II smooth muscle-dependent dilation response and each of the following 6-month outcomes: NOR (p=0.5), NOL (p=0.96) and TOR (p=0.8). These data suggest lack of association between cognitive function and arterial function following angiotensin-II exposure. Arterial dilator responses to acetylcholine and DETA NONOate were inversely related to CBV in lateral hippocampus (Figure 6B-D, Supplement Table 3).

**Figure 6.**
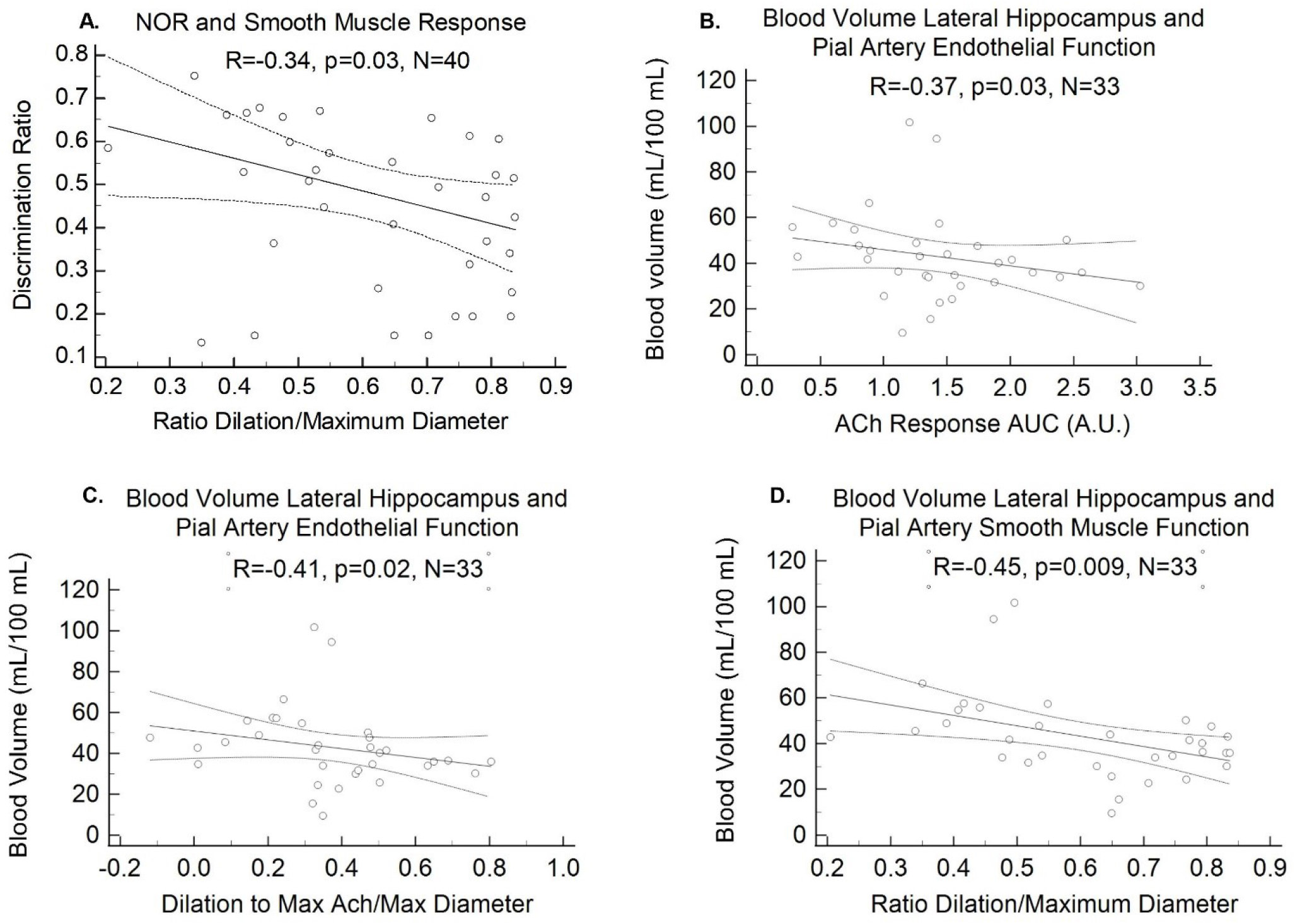
Pial arterial function and cognitive function/regional resting blood volume relationships. A. There is inverse relationship between 6-month novel object recognition score and dilator response to DETA NONOate (smooth muscle function). B-D. Resting pial arterial dilator response to acetylcholine (AUC and maximum acetylcholine dose) and DET NONOate are also inversely related to in vivo resting blood volume in lateral hippocampus

### Acute righting reflex recovery and cognitive, imaging and vasoreactivity outcomes

In brain-injured rats, correlation analyses showed no significant correlation between righting reflex recovery time post-injury with the following outcome measures: 3- and 6-month NOR, NOL, TOR, 6-month regional and global CBF and regional CBV, and 6-month baseline dilator response to acetylcholine and DETA-NONOate, and change in dilator response to acetylcholine or DETA-NONOate following exposure to angiotensin II, high glucose or Aβ42.

## DISCUSSION

The findings demonstrate novel preclinical evidence that mild TBI from a midline fluid percussion injury^21^ results in persistent/chronic 3- and 6-month cognitive-behavioral impairment (NOR). It also shows for the first time that we know of that there are significant associations between regional cerebral blood flow in the lateral and medial hippocampus and S1BF with cognitive function following mTBI. The 6-month resting *in vivo* regional CBF and CBV and *ex vivo* baseline cerebral arterial myogenic tone, endothelial and smooth muscle function did not differ between mTBI and sham rats, but the groups differed in smooth muscle response following exposure to angiotensin II. The results provide evidence of chronic adverse consequences of mTBI, consistent with human epidemiologic cross-sectional observations.^4^

TBI is a main cause of death and disability in the US in people younger than 35 years.^4^ Case control and epidemiologic studies indicate that individuals with TBI history have a higher risk of developing dementia.^11, 12^ Preclinical studies in severe TBI show late (1-year) development of cognitive impairment following injury,^16–19^ but empiric evidence as regards chronic consequence of mTBI is lacking. In the midline fluid percussion model used in this study, there are immediate transient deficits, which transition to late-onset morbidities even in the absence of gross histopathology,^21^ making this a useful model of mild TBI that resulted in chronic impairments. Our results show that even mTBI leads to chronic persistent cognitive dysfunction present at 3 months and sustained at 6 months post-injury. This is consistent with clinical observations that even mild head injury was found to be a predisposing factor for some cases of Alzheimer’s disease.^14^ Our findings therefore validate the use of this injury model to explore mechanisms by which mild TBI leads to late dementia.

Disturbed cerebrovascular function was observed in humans following TBI in the acute (≤1 day) and subacute (1 week) period ^15, 35^ with inverse relationship between CBF and cognitive function.^15^ Vascular injury is known to contribute to severe TBI neuropathology with ischemic brain damage being found on autopsy in >90% of acute TBI mortalities.^6^ Severe TBI acutely resulted in reduced local CBF and neurovascular uncoupling^36^ with endothelial dysfunction.^37^ Using the same fluid percussion injury model, we previously showed regional morphologic cerebrovascular changes (increased average cerebral arterial vessel volume and surface area) 7 days post-mTBI likely representing an acute response to mechanical forces of injury.^23^ In contrast to empiric data in the acute setting, data on cerebrovascular flow/function and their relationship to cognitive function in the chronic setting post-mTBI are lacking. Our findings show little difference in global and regional CBF and CBV 6 months post-mTBI in the rat model, similar to observations in human concussion patients where CBF normalizes at 1-month post-injury.^15^ These findings suggest that the vascular perturbations observed previously in the acute setting following mTBI recover in the chronic setting. At 6 months we found a direct relationship between regional CBF (lateral and medial hippocampus and S1BF) and cognitive function (NOR and NOL), similar to the association between CBF and neuropsychiatric function in the acute setting following concussion in humans.^15^ The lack of difference in resting CBF and CBV at 6 months between mTBI and sham rats does not necessarily rule out the causative role of vascular perturbations in chronic cognitive dysfunction in mTBI. The persistence of cognitive dysfunction, but not resting CBF and CBV abnormality, suggest that the vascular influence modulating chronic cognitive function could be predominant early in the injury process. Additionally, CBF and CBV were measured in a basal state devoid of functional challenges that could uncover latent vasoreactive deficits.

Results show no difference in pial cerebral arterial myogenic tone and resting endothelium-dependent and smooth muscle-dependent function between mTBI and sham. Persistent vasculopathy, however, is suggested by our finding of the variant dilator responses of mTBI cerebral arteries following exposure to angiotensin II when compared to sham arteries. Sham arteries exposed to angiotensin II demonstrate reduced dilation to DETA NONOate and acetylcholine compared to pre-exposure responses whereas mTBI arteries showed no difference, demonstrating reduced vasoconstrictor response to angiotensin II in mTBI arteries. Angiotensin II is an important regulatory peptide mediating vascular oxidative stress, inflammation and vasoconstriction via action on angiotensin II type 1 receptor in vascular smooth muscle cells.^38^ Acute short-term exposure of epineural arterioles to angiotensin II resulted in decreased blood flow in normal peripheral nerves^39^ and altered vasoreactivity response to angiotensin II was shown in diabetic animals when compared to normal controls.^39–41^ The lack of difference in baseline dilator response to acetylcholine and DETA NONOate, but with differential response following exposure to angiotensin II between mTBI and sham parallels, but is opposite to, observations in another neurodegenerative condition. Isolated gluteal resistance arteries from patients with cerebral autosomal dominant arteriopathy with subcortical infarcts and leukoencelopathy (CADASIL) showed similar vasoreactive responses to acetylcholine and spermine-NONOate as controls, but CADASIL arteries showed greater vasoconstrictor response to angiotensin II than controls.^42^ Data on cerebral arterial response was not available for this cohort which may be relevant as effects of angiotensin II were shown to vary in different arterial beds.^39^ The pathophysiologic consequence of the altered arterial vasoreactivity response following angiotensin II exposure in mTBI requires further mechanistic investigation. In contrast to angiotensin II, no difference was noted between mTBI and sham following exposure to Aβ42, the amyloidogenic protein implicated in Alzheimer’s disease that also induces cerebral arterial endothelial dysfunction,^33^ or to high glucose, the metabolic abnormality in diabetes mellitus that also causes acute endothelial dysfunction.^34^

Our results showed significant correlation between 6 month NOR and baseline dilator response to DETA NONOate, suggesting the association between cognitive function and cerebral arterial smooth muscle function. The lack of correlation between measures of cognitive function and change in dilator response following angiotensin II exposure suggests this vascular perturbation could not explain the chronic cognitive dysfunction observed in this model.

Among the brain regions measured, only the lateral hippocampus resting CBV was related (inversely) to ex-vivo pial cerebral artery endothelial and smooth muscle function. Whether this relationship represents enhanced influence by and/or greater vulnerability in this brain region to perturbations in large cerebral arterial function is unknown and needs to be explored further. A hippocampal subfield volumetry study in cognitively normal and mild cognitive impairment patients showed region-specific vulnerability of hippocampal subfields to vascular injury with differential hippocampal subfield atrophy patterns seen between MCI from vascular disease versus non-vascular causes.^43^

Our study has several limitations. We only studied male rats, so chronic mTBI effects on female rats need to be studied in the future. We show novel cerebrovascular and cognitive function data 6 months following mTBI but lack data on early post-injury cerebrovascular function that could clarify the role of early vascular perturbation in the persistent cognitive impairment observed. We showed *in vivo* resting CBF and CBV but lack data on *in vivo* cerebrovascular reserve that could be shown by performing hypercapnic responsiveness or assessing neurovascular coupling. *Ex vivo* results following exposure of pial arteries to angiotensin II uncovered differential responsiveness between mTBI and sham rats suggest the need for *in vivo* assessment not only of resting but also post-stress cerebrovascular function. Based on the associational relationships uncovered by this study, future efforts can follow to establish causal mechanisms using interventions that modulate vascular conditions to establish the role of vascular dysfunction in chronic TBI-mediated cognitive impairment. The subtle magnitude of the chronic neurologic and vascular changes observed may reflect the mild TBI nature of the injury model and the intervening endogenous repair mechanisms, yet the chronic pathophysiology shows similarity to cross-sectional observations in human mild TBI,^10, 14^ enhancing the utility of this model and the neurovascular findings. Lastly, there currently exists no consensus classification schema for TBI severity,^44^ and our injury model remains consistent with criteria for mild TBI using either Department of Defense or Veterans Affairs classification schemes;^10^ future consensus classification may alter the designation of (or be informed by) our model.

In conclusion, mTBI resulted in chronic 3- and 6-month cognitive dysfunction and altered cerebral arterial vasoreactivity response following exposure to angiotensin II, without a change in 6-month resting CBF, CBV or baseline endothelial or smooth muscle dependent function. The results demonstrate persistent pathophysiologic consequences of mTBI that could translate to human exposure to mild TBI.

## Supporting information

There are three tables of supplementary material for this paper.

## ACKNOWLEDGEMENTS

We thank Kyle Offenbacher and Karen D’Souza for their assistance with experimental procedures. We thank Bret Tallent, Gail Farrell, and the Arizona Veterans Research and Education Foundation for their assistance.

## AUTHOR CONFIRMATION STATEMENT

RQM, JL and CQ made substantial contribution to the concept and design. DRG, RQM, LML, AF, LCB, GT and CCQ wrote the manuscript draft and JL, PDR and RG critically revised the manuscript. AF, ST, NK, HE, DM reviewed and approved the manuscript. DRG, LML, CY, AF, ST, NK, GT, HE acquired the data, RQM, JL, CCQ, LCB and AF analyzed and interpreted the data. All authors reviewed and approved the version of the manuscript submitted.

## DATA SHARING

The data are available upon reasonable request.

## SUPPLEMENTAL MATERIAL

There are three tables of supplementary material for this paper.

