## Supplementary material for "Chronic Cognitive and Cerebrovascular Function Following Mild Traumatic Brain Injury in Rats": There are three tables of supplementary material for this paper.

Supplement Table 1. Breakdown of animal outcome measures.

| | Sham | | mTBI | |
|---|---|---|---|---|
| | 3 months | 6 months | 3 months | 6 months |
| Cognitive Function Assessment | 21 | 21 | 20 | 20 |
| Imputed Score: NOR | 1 | 3 | 2 | 4 |
| Imputed Score: NOL | 1 | 3 | 2 | 4 |
| Imputed Score: TOR | 3 | 3 | 1 | 3 |
| MRI/Vasoreactivity | | 17 | | 15 |

Supplement Table 2. Correlation Analyses Among Cognitive Function and MRI Outcome Measures

| Imaging Parameter | NOR (6 months) | NOL (6 months) | TOR (6 months) |
|---|---|---|---|
| CBF Global | R=0.17<br>p=0.35 | R=0.11<br>p=0.55 | R=0.01<br>p=0.94 |
| CBF Lateral Hippocampus | **R=0.54**<br>**p=0.01** | R=0.23<br>p=0.32 | R=0.24<br>p=0.30 |
| CBF Medial Hippocampus | **R=0.57**<br>**p=0.007** | **R=0.54**<br>**p=0.01** | R=0.18<br>p=0.44 |
| CBF S1BF | **R=0.62**<br>**p=0.003** | R=0.23<br>p=0.31 | R=0.12<br>p=0.62 |
| CBV Lateral Hippocampus | R=0.22<br>p=0.22 | R=-0.07<br>p=0.7 | R=0.10<br>p=0.58 |
| CBV Medial Hippocampus | R=-0.18<br>p=0.32 | R=-0.23<br>p=0.21 | R=-0.22<br>p=0.23 |
| CBV S1BF | R=0.18<br>p=0.33 | R=0.07<br>p=0.69 | R=-0.17<br>p=0.35 |

CBF- cerebral blood flow, CBV-cerebral blood volume, NOL- novel object location, NOR-novel object recognition, TOR- temporal order object recognition, S1BF-primary somatosensory barrel cortex

Supplement Table 3. Correlation Analyses Among Vasoreactivity Outcomes and Cognitive/MRI Outcomes

| Outcomes | Max. Ach Dilation | Dilation to DETA-NONOate |
|---|---|---|
| NOR (6 months) | R=-0.26<br>p=0.10 | **R=-0.34**<br>**p=0.03** |
| NOL (6 months) | R=-0.05<br>p=0.75 | R=-0.13<br>p=0.44 |
| TOR (6 months) | R=0.15<br>p=0.35 | R=0.02<br>p=0.88 |
| CBF Global | R=-0.08<br>p=0.68 | R=-0.10<br>p=0.57 |
| CBF Lateral Hippocampus | R=-0.24<br>p=0.29 | R=-0.1<br>p=0.66 |
| CBF Medial Hippocampus | R=-0.18<br>p=0.42 | R=-0.30<br>p=0.18 |
| CBF S1BF | R=-0.27<br>p=0.24 | R=-0.35<br>p=0.12 |
| CBF Thalamus | R=0.1<br>p=0.68 | R=-0.30<br>p=0.18 |

| | | |
|---|---|---|
| BV Lateral Hippocampus | **R=-0.41** **p=0.02** | **R=-0.45** **p=0.009** |
| BV Medial Hippocampus | R=-0.165 p=0.36 | R=-0.11 p=0.55 |
| BV S1BF | R=-0.23 p=0.2 | R=-0.23 p=0.19 |
| BV Thalamus | R=-0.05 p=0.78 | R=-0.12 p=0.50 |

CBF- cerebral blood flow, CBV-cerebral blood volume, Dilation to DETA-NONOate- baseline dilation response to $10^{-4}$M diethylenetriamine NONOate, Max Ach dilation- baseline dilation response to acetylcholine $10^{-4}$M, NOL- novel object location, NOR-novel object recognition, TOR-temporal order object recognition, S1BF-primary somatosensory barrel cortex
